# Accelerating Strain Engineering using Desorption Electrospray Ionization-Imaging Mass Spectrometry and Untargeted Molecular Analysis of Intact Microbial Colonies

**DOI:** 10.1101/2021.04.01.438078

**Authors:** Berkley M. Ellis, Piyoosh Babele, Jody C. May, Carl H. Johnson, Brian F. Pfleger, Jamey D. Young, John A. McLean

**Author notes:** Correspondence: John A. McLean.

## Abstract

**Progress in the fields of genomic and biologic sciences has yielded microbial bioprocesses for the advanced production of chemicals. While biomanufacturing has the potential to address global demands for renewable fuels and chemicals, engineering microbial cell factories that can compete with synthetic chemical processes remains a challenge. Optimizing strains for enhanced chemical production is no longer limited by reading and writing DNA, rather it is impeded by the lack of high-throughput platforms for characterizing the metabolic phenotypes resulting from specific gene editing events. To address this issue, we have developed a desorption electrospray ionization- imaging mass spectrometry (DESI-IMS) screening assay that is conducive to both multiplexed sampling and untargeted analyses. This technology bridges the gap between genomic and metabolomic timescales by simultaneously characterizing the chemical output of various engineered *Escherichia coli* strains rapidly and directly under ambient conditions. The developed method was used to phenotype four *E. coli* strains on the basis of measured metabolomes, which were validated via PCR genotyping. Untargeted DESI-IMS phenotyping suggests multiple strategies for future engineering which include: (i) relative amounts of specific biosynthetic products, (ii) identification of secondary products, and (iii) the metabolome of engineered organisms. In sum, we present a workflow to accelerate strain engineering by providing rapid, untargeted, and multiplexed analyses of microbial metabolic phenotypes.**

---

**E**ngineering microorganisms for the renewable manufacturing of chemicals is becoming increasingly viable given major advancements in genome and biological sciences. Biosynthetic production of commodity chemicals as an alternate path to traditional chemical synthesis has the potential to address global challenges across medical, environmental, energy, and economic sectors.^1–3^ However, engineering microorganisms, such as bacteria, with the efficiency and low cost of commercial chemical syntheses remains a significant challenge. Despite the ability to rapidly alter and edit bacterial genomes, directing these alterations to elicit specific biological changes is hindered by the rate at which phenotypic outcomes can be measured from individual gene edits.^2,4–6^ In part, this is due to a fundamental mismatch in the throughput with which genetic editing can be performed versus how quickly the metabolic consequences can be determined. Indeed, comprehensive chemical characterization of engineered strains is considered one of the grand challenges in synthetic biology.^2,7,8^ Thus, increasing the throughput and molecular breadth of analytical readouts provides opportunities to accelerate metabolic engineering and ultimately promote commercial feasibility of microbial cell factories.

Contemporary synthetic biology workflows for optimizing microbial biosynthesis are rooted in functional genomics, directed evolution, and metabolic flux studies all of which depend on chemical measurements. To effectively support these applications, the analytical method utilized must be rapid, minimally invasive to microbial colonies, and broadly amenable to many molecular classes. Label free sampling provides a more rapid and robust annotation of the microorganism’s chemical productivity by alleviating the need for additional engineering steps encoding sensors, or probes, and further perturbing the system. Contemporary fluorescent spectroscopy methods are rapid and feasible if the desired product is naturally fluorescent, however, the majority of biomolecules do not intrinsically fluoresce, which limits the chemistries that can be targeted.^6,9^ Techniques such as riboswitches, RNA aptamers, ligand responsive transcription factors, or coupled enzyme reactions can also be used for rapid screening of a wider variety of chemistries, but at the cost of additional engineering steps that may confound readout of the biological system being investigated.^6^ Furthermore, the methods reported thus far have been tailored towards specific products that are not yet fully generalizable.^6^ Mass spectrometry-based techniques address many of these challenges and limitations by simultaneously measuring the intrinsic property of mass-to-charge ratio (m/z) of a large breadth of molecular species at rates commensurate with high throughput analyses.^8^

Most mass spectrometry (MS) methods to investigate synthetic biology use gas chromatography-mass spectrometry (GC-MS) or liquid chromatography-mass spectrometry (LC-MS) to target specific endpoint molecules arising from microbial biosynthesis.^10,11^ Both GC-MS and LC-MS typically require several steps of sample handling prior to MS analysis, including analyte extraction, liquid sample preparation procedures, and, in some instances, chemical derivatization to enhance sensitivity for minimally volatile analytes. Furthermore, GC-MS and LC-MS methods can only analyze one sample (e.g. strain extract) at a time, and chromatographic separations necessitate several minutes to complete per sample.^10,11^ To reduce sample handling and increase throughput, a recently developed MS method using matrix assisted laser desorption/ionization (MALDI) provided rapid sampling from a 96-well plate of bacterial samples.^12^ However, this sequential MALDI strategy is not conducive to analyzing most small molecules due to matrix interferences and the need for vacuum-based sampling, which otherwise restricts the ability to measure labile and volatile analytes.^12,13^ Additionally, these chromatography and MALDI-based methods are not amenable to direct sampling of microbial colonies. Previously, Barran and coworkers described a targeted method based on desorption electrospray ionization (DESI) with MS analysis to measure biocatalysis from live bacterial colonies.^14^ Whereas this approach enabled the direct sampling of microorganisms, the targeted assay focused exclusively on tracking the conversion of specific molecules and did not examine the broader metabolome of the microorganisms. We have recently described an aligned DESI-MS strategy operated in an imaging MS (IMS) mode for rapid and reproducible analysis of microbially produced metabolites and chemicals *in situ*.^15^ Building on this approach, we have tailored this sampling method for the simultaneous analysis of many bacterial colonies including multiple strains and biosynthetic products in a spatially-resolved manner. Herein, we describe the application of DESI-IMS to untargeted metabolomic characterization of engineered microorganisms. With this technique, we obtain analytical information simultaneously for both the targeted biosynthetic products and also global metabolites detailing the broader biology of the organisms. We demonstrate the capabilities of this DESI-IMS workflow using *Escherichia coli* strains that have been genetically engineered to overproduce free fatty acids (FFA) with defined chain-lengths by expressing different thioesterase enzymes: strain TY05 overproduces C12 FFAs, strain NHL17 overproduces C8 FFAs, WT-TesA overproduces a range of C8-C14 FFAs, and control strain TY06 is genetically identical to TY05 but expresses a non-functional thioesterase.^16–19^ We highlight the advantages of untargeted acquisitions and unsupervised data analytics by characterizing secondary fatty acid products secreted by these strains. Untargeted metabolomic analyses additionally provide insight into the molecular profile of the microorganisms via variations in membrane lipid saturation, which has been shown to affect cell viability.^20^ The untargeted DESI-IMS method described in this work provides a rapid and transferable method that can be readily adapted to a wide range of biosynthetic products from fatty acids to antibiotics as well as synthetic biology strategies including directed evolution, functional genomics, and metabolic flux studies.

## Results and Discussion

### Rapid Metabolic Phenotyping of FFA Overproducing E. coli

Engineering *E. coli* fatty acid metabolism has led to the production of high-energy density, liquid transportation fuels and high-value oleochemicals from renewable feedstocks.^21,22^ With reported amounts reaching up to 70% of the theoretical yield, engineers have achieved higher production of FFA than any other downstream products.^21,22^ If these yields can approach theoretical limits, then processes may be developed to replace current petrochemical production strategies.^22^ Acyl-acyl carrier protein (ACP) thioesterases are some of the primary targets for engineering FFA production.^22,23^ These enzymes regulate the catalytic link between fatty acid biosynthesis (FAB) and a product sink, increase flux by depleting the primary regulatory signaling molecules, and control the composition and chain length of the final products.^23^ To enhance the activity of these enzymes and thus create more productive strains directed evolution or mutagenesis studies can be performed.^24,25^ In particular, screening libraries with varying modifications to acyl-ACP thioesterases can provide pivotal insight to controlling chain lengths and compositions of free fatty acids, which is a challenge within this field.^26^

A previously developed microporous membrane scaffold method was adapted to sample a mixed culture of engineered *E. coli* strains (figure 1).^15^ Briefly, microorganisms were grown on a membrane scaffold placed on top of an agar substrate, which facilitated the uptake of nutrients to support microbial growth and production, while retaining microorganisms and their excreted biosynthetic products on the scaffold surface. At desired timepoints, the membrane was removed from the agar and affixed to a glass microscope slide to provide a solid support on which DESI can be performed directly with no further sample pretreatment. The typical DESI-IMS pixel size used in these experiments was 50 × 250 μm. We tested the effect of raster rate on monocultures of the C12-producing strain TY05 using dodecanoic acid (C12:0) as a read out, and found that increasing the conventional raster rate by 50% (100 μm/s to 150 μm/s) had negligible effects on the measured amounts of dodecanoic acid detected when compared to background (supplemental figure 1). However, only using conventional parameters, we were able to completely sample 42 colonies in 356 minutes. Including the time needed to transfer the samples onto the DESI slide (ca. 10 minutes), the sample throughput was 8 minutes per microbial colony from solid-phase growth to acquired data. While MALDI-MS is capable of very fast acquisition speeds on the orders of seconds, there are additional throughput restraints imposed by colony picking, matrix application, and sample loading that are not generally reported in these workflows.^12,13^ Thus, we concluded that the 8 minutes needed to obtain the full MS images of each colony was suitable for rapid screening applications.

**Figure 1.**
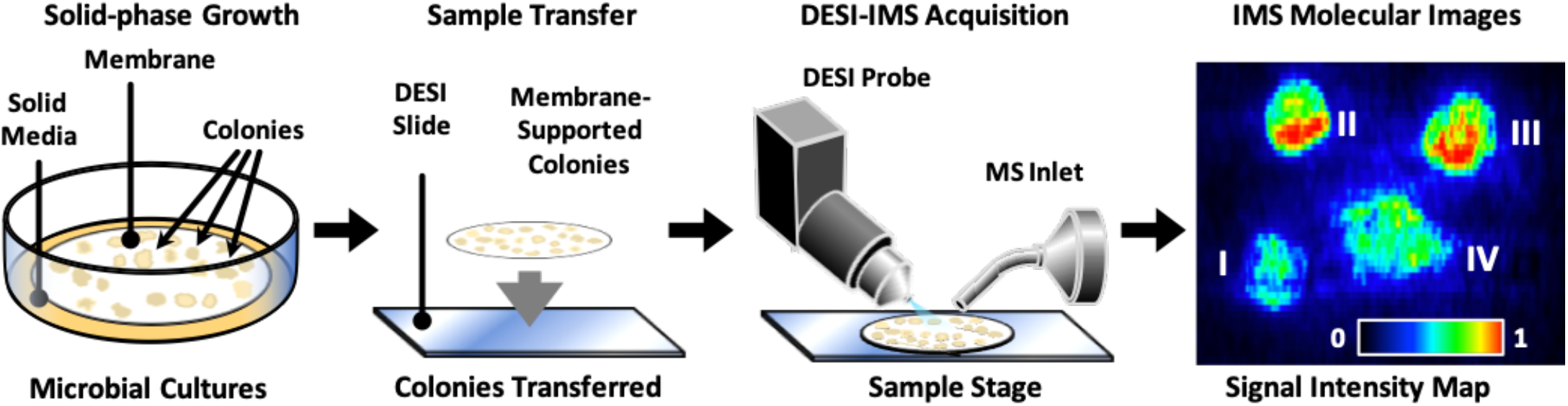
Sample preparation and acquisition workflow to profile microbial biosynthesis. Microorganisms are grown on a membrane scaffold placed on top of agar. The microorganisms and scaffold are removed from the agar and adhered to a glass slide. The DESI directly samples the microorganisms, biosynthetic products, and metabolites from the scaffold in a spatially resolved MS analysis (IMS). The resulting IMS image maps the relative amounts of different chemical products and metabolites to spatial positions (e.g., within each colony).

DESI-IMS facilitates the direct analysis of microbial colonies, while acquiring chemical information on biosynthetic products including relatively volatile analytes (e.g., short and medium chain fatty acids). Additionally, IMS is inherently a multiplexed technique through the acquisition of full MS spectra at each sampling location. We demonstrate these multiplexing and volatile analyte detection capabilities for the DESI-IMS analysis of a co-culture of two engineered *E. coli* strains TY05 and NHL17 (figure 2). In a co-culture with isopropyl-D-thiogalactopyranoside (IPTG) present to induce FFA overproduction, TY05 is expected to preferentially produce dodecanoic acid (C12:0), whereas NHL17 produce octanoic acid (C8:0).^16,17^ The spatial location of each colony was determined using an ion image for a lipid common to both strains, phosphatidylglycerol (16:0/17:1), whereas the ion images corresponding dodecanoic acid and octanoic acid were localized to specific colonies, which readily distinguished the identities of each colony (figure 2A-B). Based on the expected results of the gene edits, we predicted that the colonies that co-localized with higher intensities of dodecanoic acid are the TY05 strain and those that co-localized with accumulation of octanoic acid are the NHL17 strain.^16,17^

**Figure 2.**
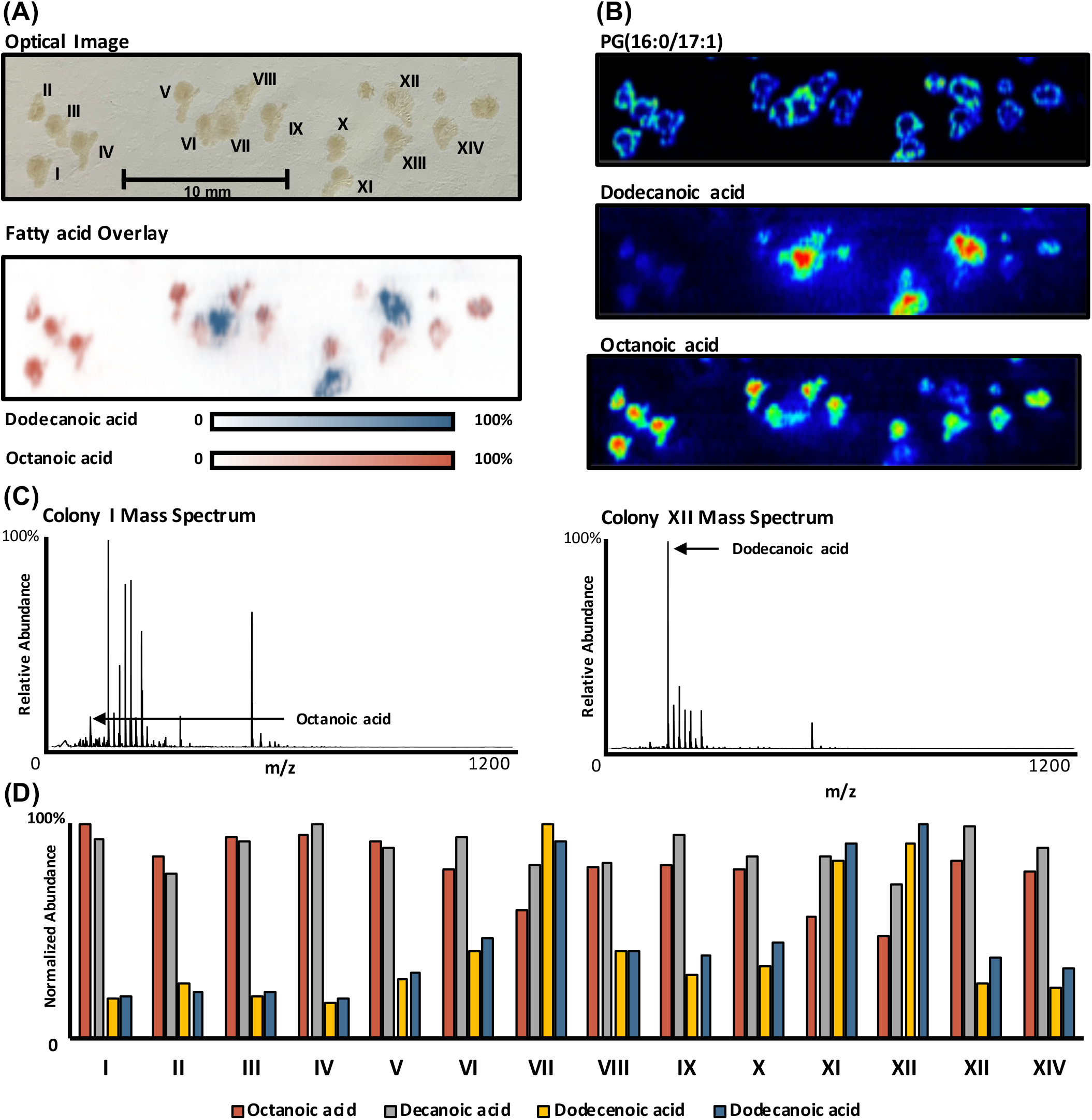
DESI-IMS data and results from a single IMS acquisition of the two engineered strains TY05 and NHL17. (A) Optical image labled with colony numbers. Fatty acid overlay demonstrates the spatial localizations of primary biosynthetic products, that correlate with strain identities. (A) Spatial mapping of a lipid common to both strains, phosphatidylglycerol (16:0/17:1), which was used to define chemical boundaries of each colony for generation of mass spectra and fatty acid intensities. Ion images of dodecanoic acid and octanoic acid highlight the relative abundance of primary biosynthetic products individually. (C) Area integrated mass spectra highlight contributions of octanoic acid and dodecanoic acid to the measured metabolomes of TY05 and NHL17 strains. (D) Normalized fatty acid abundances for each colony.

In addition to differentiating these strains through biosynthetic products, we can further characterize NHL17 and TY05 through their measured metabolomes. To demonstrate this untargeted approach, we defined colony areas as regions of interest (ROI) and integrated the MS intensities associated with Colony I and XII to generate composite mass spectra representative of each strain (figure 2C). A comparison of these two integrated mass spectra reveals distinct metabolites associated with each strain. Of note, we observe an overall higher fatty acid production from colony XII (TY05 strain) and a predominant fatty acid peak corresponding to dodecanoic acid. In the composite mass spectrum obtained from colony I (NHL17), fatty acid abundances were lower than what was observed from colony XII, which may be due to the fact that under ambient conditions octanoic acid is in the liquid phase and thus subject to greater diffusive losses than the higher chain-length FFAs. It is important to note that while fatty acids dominate the representative mass spectra, higher mass analytes (such as the PG (16:0/17:1) lipid used for colony localization) also appear at lower abundances. As per the multiplexed nature of this analysis, an ion image can be projected for any metabolite observed in the sample, allowing each bacterial strain to be phenotyped based on the totality of chemical signatures present.

To determine fatty acid abundances, we normalized the signal intensities of acquired DESI-IMS data using an internal standard (pentadecanoic acid) introduced directly to the DESI ionization solvent. This provided a readout of fatty acid (FA) ionization at each image pixel. Next, colony areas were defined as ROIs using PG (16:0/17:1) ion image to determine each colony’s chemical boundaries. Using these ROIs, total ion signals for discrete FFA mass measurements were integrated across each colony area to determine the FFA output of each colony (figure 2D). By comparing the FFA-specific output of each colony, we were able to determine that colony number XII and I are the highest producers of dodecanoic acid and octanoic acid, respectively (figure 2D).

These chemical comparisons are not limited to profiling two primary analytes. Using the same DESI-IMS data, we were able to perform a more comprehensive characterization of fatty acid biosynthesis by calculating the relative amounts of C8-C12 FFAs for each colony (figure 2D). As noted previously, dodecanoic acid is consistently present in greater relative abundances than octanoic acid. For decanoic acid, we did not observe significant changes in the intensities from colony to colony. We also find that colonies producing the highest amounts of dodecanoic acid also seem to be producing the unsaturated fatty acid dodecenoic acid (C12:1) as well (figure 2D). This may be attributed to off-target production within the TY05 strain, which suggests an additional engineering strategy for TY05. More selective production may be achieved by knocking out the desaturase enzyme(s) responsible for the conversion of C12:0 to C12:1 to prevent the formation of the C12:1 byproduct. With this untargeted approach, we highlight both the relative amounts of biosynthetic products and specific information on the FFA chain lengths and degrees of unsaturation in off-target production. This chemical information has implications in the rational design of strains but is particularly applicable to randomly generated strains in directed evolution studies. By comprehensively characterizing biosynthesized products, the potential for discovering novel and favorable mutations is more feasible than profiling a specific product. Thus, our DESI-IMS platform can support the rapid development and discovery of strains with enhanced productivity.

### Untargeted Molecular Phenotyping via Unsupervised Segmentation

Whereas the FFAs are of immediate interest for this study, the vast majority of molecular species measured using untargeted DESI-IMS acquisitions are not free fatty acid products. These additional metabolites can provide further insights into biosynthetic processes such as product/precursor levels, product sinks, and off-target production, the latter of which has implications for discovering additional commercially relevant secondary products or wasteful byproduct pathways that reduce product yield. Furthermore, the effects of genetic edits on the growth and viability of the engineered organisms can be indicated by accumulation or depletion of metabolites outside of the primary biosynthesis pathways. This includes, but is not limited to, cellular redox through redox pairs, levels of amino acids and other central metabolites, and degrees of membrane lipid saturation. Comprehensive metabolic characterization (including biosynthetic products) of engineered organisms can identify future editing targets and strategies to enhance production without additional experiments. To facilitate the investigation of all measured molecular features within these DESI-IMS experiments, we developed a spatially oriented untargeted metabolomics data analysis workflow using unsupervised segmentation.

Unsupervised segmentation is a computational process that groups IMS pixels into chemically similar regions using measured mass spectral similarities. The end result of this process is an image with spatially oriented groups (segments) that have discrete molecular signatures. The chemical fingerprints of each segment are calculated on the basis of a significance value determined by a t-test comparing the intensity of a feature within a segment area to the rest of the IMS image.^27,28^ Thus, each feature associated with a segment is significantly upregulated within that region and also statistically absent from the remainder of the IMS image. There are many fundamental advantages to this untargeted analytics approach as the areas and even phenotypes of the microbial colonies are defined unbiasedly. This minimally-biased approach is advantageous in that it reduces the user input for analysis, removes error associated with manually defining colony areas and reduces overall analysis time. Moreover, the segments generated via discrete molecular profiles account for all of the measured features within a DESI-IMS experiment, which lead to a more robust assignment of phenotypes. This type of analysis includes any biomarkers that may be expressed by strains as a result of specific genetic edits. For instance, in our FFA-producing *E. coli* panel we may identify various lipid biomarkers as a result of the different alterations to fatty acid biosynthesis pathways. We compared unsupervised segmentation against other IMS data analysis methods such as ROI integration and MS profile averaging, and found no significant differences in the reproducibility and reported amounts of biosynthetic products (supplemental figure 2). Thus, unsupervised segmentation provides untargeted metabolomic phenotyping across the measured metabolome while accurately and reproducibly characterizing biosynthetic products.

We applied our developed DESI-IMS and unsupervised segmentation workflow to a mixed culture comprised of the control strain TY06 strain and FFA-overproducing strains TY05, NHL17, and WT-TesA that were collectively seeded on the microporous membrane scaffold.^16–18^ This facilitated the formation of colonies for each strain that were distributed throughout the plate area randomly and unbeknownst to the experimenter. Unsupervised segmentation separated the IMS image into 6 primary segments representing 6 distinct molecular profiles (figure 3). These 6 segments were subsequently classified into four segments representing colony areas, one segment representing the membrane scaffold, or background, and one representing the area surrounding each colony. While we anticipated only 5 segments (4 strains and 1 background), the additional segment discovered by the unsupervised method (colony margins) can provide additional insight into the growth and biosynthesis of these engineered strains. Unsupervised segmentation revealed accumulation of fatty acids in the area surrounding the colonies that were not expected based on the strain engineering strategy, namely nonadecenoic acid (C19:1), heptadecanoic acid (C17:0), and hexadecenoic acid (C16:1). We also found tetradecanoic acid (C14:0) and PG (16:0/15:1) to be in significant abundance within these boundary regions. This segment and the contributing metabolites may represent biomarkers of colony growth and expansion, byproducts stemming from lipid biosynthesis, or even diffusion of chemicals away from the colonies. More notably, this additional segment demonstrates the sensitivity of the DESI-IMS sampling method and the utility of unsupervised segmentation to discover unanticipated molecular signatures.

**Figure 3.**
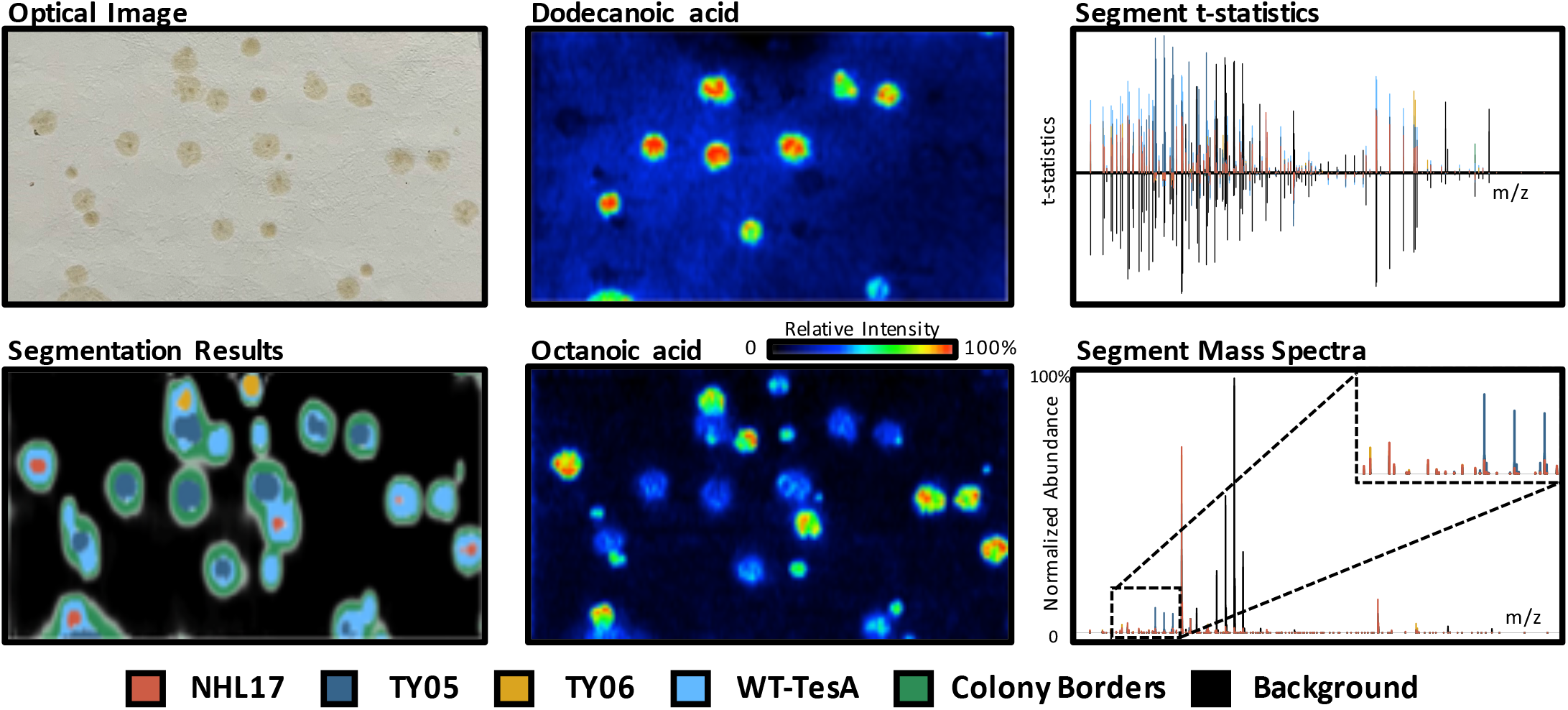
Unsupervised segmentation unbiasedly determines molecular phenotypes of engineered microorganisms on the basis of measured metabolomes. Segment statistics and mass spectra highlight the primary features contributing to each strain segments. The segmentation results show the unique phenotypes of the four strains analyzed in this experiment. The dark blue segment represents the dodecanoic acid producing TY05 strain and red corresponds to NHL17 engineered for octanoic acid production. The yellow and light blue segments represent the control TY06 strain and the broad FFA producing WT-TesA strain. We validated the identification on the basis of measured metabolomes

The four discrete molecular profiles associated with the bacterial colonies represent four unique metabolic phenotypes. To determine which strain these phenotypes corresponded to, we used t-statistic values and average mass spectrum for each segment (figure 3). With prior knowledge of the anticipated FFA production of each strain, we were able to infer strain identities using the measured FFA abundances. For instance, dodecanoic acid had the highest t-statistic value in the dark blue segment, which ultimately led to the associated colonies being assigned as the C12-producer TY05 (figure 3).^17^ The highest average intensity of dodecanoic acid observed across all MS data was within this segment, which further supported this assignment. Additional analyses were required to distinguish the strains associated with octanoic acid. The WT-TesA strain overproduces a range of FFA chain-length distributions including C8:0, which is also the primary product of NHL17. ^16,18^ Due to the labile nature of octanoic acid yielding lower signal intensities in the DESI-IMS analysis, both the red and light blue segments had similar t-statistic values (figure 4). However, when we considered the average metabolite intensities for octanoic acid in the segments, we found that the red segment had nearly twice the amount of octanoic acid per pixel. This led us to tentatively assign the NHL17 strain to the red segment. To solidify our assignments, we characterized the abundances of fatty acids ranging from C8-C14 within the unassigned light blue and red segments (supplemental figure 3). When comparing the segment fatty acid profiles, we found that the light blue segment exhibited broader FFA production, particularly we found elevated amounts of dodecanoic acid (C12:0), tetradecenoic acid (C14:1), and tetradecanoic acid (C14:0). Further, we observed higher accumulation of octanoic acid in the red segment (supplemental figure 3). This confirmed our initial assignment of the NHL17 strain to the red segment and also designated the light blue segment as the WT-TesA strain. By process of elimination, the yellow segment was assigned as the TY06 strain. Additionally, the segment fatty acid profiles, t-statistics, and average metabolite abundances all indicated that this segment correlated to the minimal fatty acid production, supporting our identification. Importantly, all of these strain assignments were validated using a PCR assay (supplemental figure 4). Thus, DESI-IMS and unsupervised segmentation corroborates established methods for identifying strains and effectively determines strain identities on the basis of the measured surface metabolome. Notably, this DESI-IMS phenotyping of specific gene edits is performed in a rapid, untargeted, and multiplexed manner.

**Figure 4.**
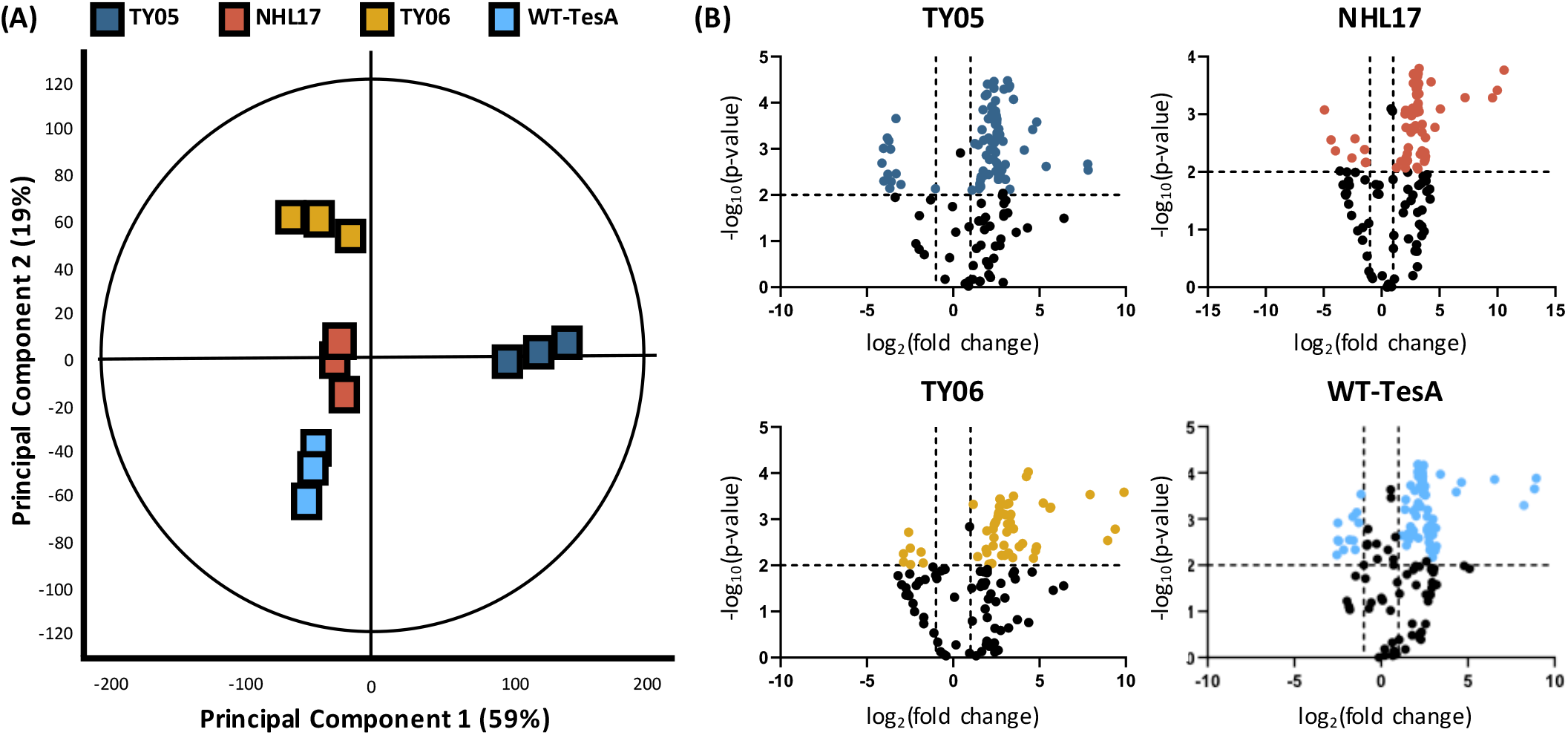
(A) PCA demonstrating the overall metabolic distinctions associated with the NHL17, TY05, TY06, and WT-TesA segments. (B) Volcano plots comparing observed features from each strain compared to the background segment. The highlighted points represent the significant (p-value ≥0.05, |fold change| ≥2) molecular features which contribute to each strain’s unique phenotypes.

### Unsupervised segmentation of DESI-IMS measurements elucidates the underlying biology of engineered microorganisms

To demonstrate the numerous features that are changing from segment to segment, we used traditional metabolomic statistical methods including principal component analysis (PCA) and volcano plot projections (figure 4). The mass spectral feature intensities associated with each strain segment were integrated to perform these statistical analyses in a spatially resolved manner. The PCA plot demonstrates that the sample replicates grouped together and partitioned from one another across two principal components (PC1 and PC2), indicating these molecular profiles were unique to each bacterial strain (figure 4A). Examination of the loadings plot corroborates the differences measured by segmentation between these sample types (supplemental figure 5). Specifically, we observe that dodecanoic acid is one of the primary features in PC1, which explains the separation between TY05 and the remaining sample types. We also found that octanoic acid was one of the distinguishing metabolites in PC2, which particularly differentiates NHL17 from WT-TesA and TY06 strains. The primary features associated with TY06 are amino acids, particularly glutamic acid, on the basis of accurate mass measurement. We also observe the unidentified features m/z 615.17 and 154.95 that contribute to the differences in TY06 and WT-TesA sample types, respectively. These and other unidentified features represent the potential for discovery using an untargeted metabolomics approach and may provide avenues for future engineering strategies for these strains.

While PCA can assess the broad scale molecular differences across different engineered strains, volcano plots highlight the specific compounds that contribute to these unique molecular profiles. The volcano plots shown in figure 4B demonstrate the statistical significance of the features within a segment as compared to the membrane background and further exhibit the number of features changing within each region. For instance, we found that NHL17 had 46 statistically upregulated features against the background compared to 59 of TY05. Further investigation of the most significant features (p-value ≥ 0.05, |fold change| ≥ 2) originating from NHL17 reveal that the most changing features correspond to varying ion species of octanoic acid. However, for TY05, we found that multiple fatty acids were significantly upregulated, particularly tetradecanoic acid (C14:0) and tetradecenoic acid (C14:1), which corroborates previous work characterizing TY05 FFA production.^17^ We also observed this off-target production of TY05 in the segment fatty acid profiles that were used to distinguish WT-TesA and NHL17 (supplemental figure 3). Within our DESI-IMS analyses, we find that dodecanoic acid (C12) represents 50% of TY05 fatty acid products within the C_8_-C_14_ range followed by tetradecanoic acid (C14) at 38%. The remaining significant fatty acid products were unsaturated fatty acids, dodecenoic acid (C12:1) and tetradecenoic acid (C14:1), contributing 2% and 6%, respectively. Detailing the composition of these biosynthesized fatty acids has critical implications in screening both directed evolution studies and rationally designed strain libraries to tailor biosynthesis towards a specific product. Further, we observe higher specificity of CpFatB1 in NHL17 than BTE in TY05.

In additional targeted analyses, we were able to putatively identify 6 lipid species that were found to be significant within the unsupervised segmentation and volcano plots analysis. We identified these lipids by incorporating tandem mass spectrometry experiments in our DESI-IMS workflow (i.e., DESI-MS/MS) and corroborating the tentative identifications via accurate mass measurement with the fragmentation results (supplemental figure 6). Of the lipids identified with DESI-MS/MS, unsaturated lipid species were found to be statistically significant in the TY05 segment compared to NHL17, WT-TesA, and TY06. In particular, we observed upregulation of phosphatidylglycerol (18:1/18:1), phosphatidylethanolamine (PE) (18:1/18:1), and PE (18:1/17:1). The increased amounts of these lipid species may contribute to membrane stress and affect the overall health and viability of the organism.^20^ It has been previously shown that *E. coli* engineered for medium-chain-length free fatty acid production has reduced viability and a loss of inner membrane integrity in previous work.^20^ The measured changes in lipid abundances via DESI-IMS that were revealed using unsupervised segmentation and volcano plots significance analysis specify downstream metabolic outcomes of genetic engineering that may be targeted to increase the overall growth and production of this particular strain.

## Conclusion

In this work, we demonstrate the utility of the presented DESI-IMS and spatially-resolved metabolomic analytics to support synthetic biology approaches. Direct analysis of microbial colonies is facilitated using the microporous membrane scaffold method, which reduces sample handling to a single step procedure (scaffold transfer to the DESI slide). To demonstrate the utility of the untargeted approach, we successfully identify four bacterial strains in a mixed culture on the basis of their measured metabolomes using a novel unsupervised segmentation analytic approach. Additionally, by measuring the full complement of biosynthesized fatty acids produced by these strains, we provide information on the specificity of engineered production not typically acquired using endpoint analysis. The DESI-IMS results demonstrate that a single, untargeted IMS acquisition can provide analytical information regarding: (i) the highest producer of a specific biosynthetic product, (ii) any secondary and off-target production, and (iii) the molecular profile of engineered organisms. This information has the promise to close the gap between genomic timescales and the analytical methods that assess them by providing various avenues for future engineering within a single multiplexed assay.

While we highlight the value of this DESI-IMS workflow for phenotyping engineered strains, we note that this approach is inherently a surface technique that measures excreted metabolites and molecules associated with the cell membranes. Thus, as implemented, DESI-IMS cannot readily provide readouts on intracellular metabolites as can be achieved from LC-MS and GC-MS, although including an extraction step prior to DESI analysis could provide intracellular metabolomic information. DESI-IMS is also a solid-phase screening method that can be used to select candidate strains to be further characterized in liquid culture, in which most characterizations of fermentation take place. The effects of solid-phase growth, fermentation, or co-culturing strains on biosynthetic production are not fully known. Although, co-culturing has been shown to bolster chemical production in *E. coli*, which could be examined using this method in future studies.^29^

The combination of DESI-IMS sampling and unsupervised segmentation have applications outside of the model system of FFA producing *E. coli*. DESI-IMS is amenable to a wide variety of small molecules (e.g., lipids, antibiotics, amino acids, etc.) many of which are not readily analyzed in vacuum-based ionization techniques. Further, DESI-IMS has been used to measure chemicals produced from a wide variety of microorganisms including gram-positive bacteria and fungi. ^30–32^ In sum, we present a method that rapidly and comprehensively annotates the metabolome of engineered microorganisms with the potential to broadly accelerate synthetic biology workflows.

## Materials and Methods

### Strains and culture conditions

Engineering strategies for the TY05, TY06, NHL17, and WT-TesA can be found in the respective publications.^16–18^ Briefly, TY05 (K-12 MG1655 ΔfadD::trcBTE, ΔfadE::trcBTE, ΔfadAB::trcBTE) was engineered to preferentially produce dodecanoic acid (C12:0) by inserting three copies of a codon-optimized acyl-acyl carrier protein thioesterase from *Umbellularia californica* (BTE) under the control of an isopropyl-β-D-thiogalactopyranoside (IPTG)-inducible promoter. These changes were integrated into chromosomal loci of B-oxidation genes (fadD, fadE, and fadAB).^17^ The same gene deletions were also made in a negative-control strain (TY06) containing three copies of BTE with an active-site mutation (BTE-H204A) that renders the protein nonfunctional.^17^ NHL17 (K-12 MG1655 ΔaraBAD ΔfadD::trcCpFatB1.2-M4-287) was augmented for enhanced octanoic acid (C8:0) production by inserting a single chromosomal copy of an optimized thioesterase under IPTG induction.^16^ The WT-TesA strain expresses a native thioesterase (TesA), which produces fatty acids with a range of chain-length distributions from C8-C14.^18^ For DESI-IMS analysis of co-cultures, single colonies of each strain were inoculated from freezer stocks into 3 mL of Luria-Bertani (LB) and incubated overnight at 37°C with shaking. An equal volume (500 µL each) of each strain culture with equal OD600 of 0.3 were mixed thoroughly in a sterile tube. Mixed culture was serially diluted 3-fold using sterile LB media. 100 µL of mixed culture was plated on the microporous membrane scaffold placed on the top of LB agar Petri dish with isopropyl β-D-1-thiogalactopyranoside (IPTG) (1 mM final concentration). All plates were incubated at 37 °C for 30 h for FFA production.

### Sample preparation and acquisitions

Membranes used for the microporous membrane scaffold method were 0.45 µM pore size and 47 mm diameter (Sigma-Aldrich). Prior to sampling, the membrane was removed from the medium, dried for 5 minutes, and adhered to a glass slide using double-sided scotch tape.DESI-IMS and DESI-MS/MS acquisitions were performed using a Waters Synapt G2S High Definition Mass Spectrometer (Waters Corporation, Manchester, UK) with a Waters x-y directional stage and DESI source described by *Tilner et al*.^33^ The system was mass calibrated with sodium formate salt clusters to a 95% confidence band and root mean square (RMS) residual mass ≤ 0.5 ppm. This resulted in experimental mass accuracies of generally 2-5 ppm and mass resolving powers circa 11,000. Optimized DESI source conditions were found to be a –5 kv capillary voltage, 110 ^°^ C desolvation temperature, 0.5 mPa N_2_ gas flow, and a cone voltage of 40 V. The sprayer was set at a 70^°^ angle, with the x, y, z, settings at −2, +2, and +2.75, respectively. Ionization solvent was comprised of 90/10 acetonitrile/water (Optima grade, Fisher Scientific) solution with 0.5% NH_4_OH with 0.2 ng/µL leucine-enkephalin for lock mass purposes and 0.2 ng/µL pentadecanoic acid for spatial normalization. Optical images were taken using a 12-megapixel camera. Imaging acquisitions were prepared using the HDImaging software. All images were acquired using a 50 x 250 µm pixel size with a raster rate of 100 µm/s. All analyses were performed in negative ion mode in the mass range of m/z 50-1200. DESI-MS/MS experiments were performed from *E. coli* colonies while selectively fragmenting a single ion of interest for identification purposes.

### Data processing and analysis

All imaging files were processed using the HDImaging software. The 1,000 most abundant features from each experiment were investigated. Mass measurements for the features were lock mass adjusted in 2-minute intervals to the leucine enkephalin internal standard. During this processing step an imaging text file was created by the HDImaging program. This text file was imported into R and Cardinal MSI.^27,28^ Feature intensities were normalized to the TIC and subsequently spatially normalized to the pentadecanoic acid (C15:0) internal standard intensity for each pixel to account for substrate-dependent ionization, or differential ionization in regards to the various areas and surfaces within the IMS experiment. This is a pivotal step in the normalization process as there is considerable fatty acid noise within DESI-MS analyses most likely due to the ambient conditions, which can interfere with the spatial intensities of fatty acids of interest. Features were then assigned using a 5 parts-per-million (ppm) mass measurement window.

Unsupervised segmentation using spatial shrunken centroids analysis in the Cardinal R package was used to distinguish the strains unbiasedly.^27,28^ Initial segmentation trials used a higher number of families (8-10) with varying shrinkage parameters (1-3) and radii (1-3) to remove background interferences and resolve strain identities into segments. Strains were identified using the significance and average intensity of biosynthetic products and metabolites.

DESI-ion mobility-MS and DESI-MS/MS data were used to identify measured metabolites. The HDImaging program generates a temporary data file that consists of the drift time plotted against m/z. This file was able to be directly imported into Progenesis QI for metabolite identification. The imported features were normalized to the TIC, deconvoluted, and searched against the online repositories KEGG and Chemspider. Search parameters for tentative matches were made on a threshold of 10 ppm mass accuracy and 80% isotopic similarity. Tentatively identified features using this process were subsequently fragmented via DESI-MS/MS analysis of microbial colonies to achieve putative identifications.

### Confirmation of strain genotype via PCR

To confirm the genotype of selected strains replica plates were prepared by streaking selected colonies on a fresh LB media plate. The identities of the replicate colonies were validated and confirmed by colony PCR using primers listed in Supplemental Table 1. Oligonucleotide primers were purchased from Integrated DNA Technologies, Inc. (Coralville, IA).

## Acknowledgements

This work was supported by the U.S. Department of Energy, Office of Science, Biological and Environmental Research Division under award number DE-SC00019404. Financial support for aspects of this research was also provided by The National Institutes of Health (Grants NIH NIGMS R01GM092218, NIGMS R37GM067152, NCI R03CA222-452-01 and NCI 1F32GM128344-01), the U.S. Environmental Protection Agency under Assistance Agreement 83573601, and th U.S. Army Research Office and the Defense Advanced Research Projects Agency (DARPA) under Cooperative Agreement W911 NF-14-2-0022. This work has not been formally reviewed by the EPA. The views expressed in this document are solely those of the authors and do not necessarily reflect those of the funding agencies and organizations. EPA, DARPA, and the U.S. Government do not endorse any products or commercial services mentioned in this publication.

## Supplementary Material

Figure 1: DESI-IMS Raster Rate Comparison; Figure 2: Comparison of IMS Data Analysis Methods; Figure 3: Segment FFA Profiles; Figure 4: Unsupervised segmentation results highlight strain identities as validated by PCR; Figure 5: PCA Loadings Plot; Figure 6: MS/MS Spectra of Identified Lipids Supplemental Table 1: List of *E. coli* strains, genotype and primer used for their validation.

## ABBREVIATIONS

IMS: Imaging Mass Spectrometry
LC-MS: Liquid Chromatography-Mass Spectrometry
GC-MS: Gas Chromatography-Mass Spectrometry
MALDI: Matrix-Assisted Laser Desorption Ionization
DESI: Desorption Electrospray Ionization
FAB: Fatty acid Biosynthesis
PCR: Polymerase Chain Reaction
MS/MS: Tandem Mass Spectrometry
FFA: Free Fatty Acid
PCA: Principal Component Analysis
PC: Principal Component
ROI: Region of Interest
PG: Phosphatidylglycerol
PE: phosphatidylethanolamine

