## Supplementary Material for "Accelerating Strain Engineering using Desorption Electrospray Ionization-Imaging Mass Spectrometry and Untargeted Molecular Analysis of Intact Microbial Colonies"

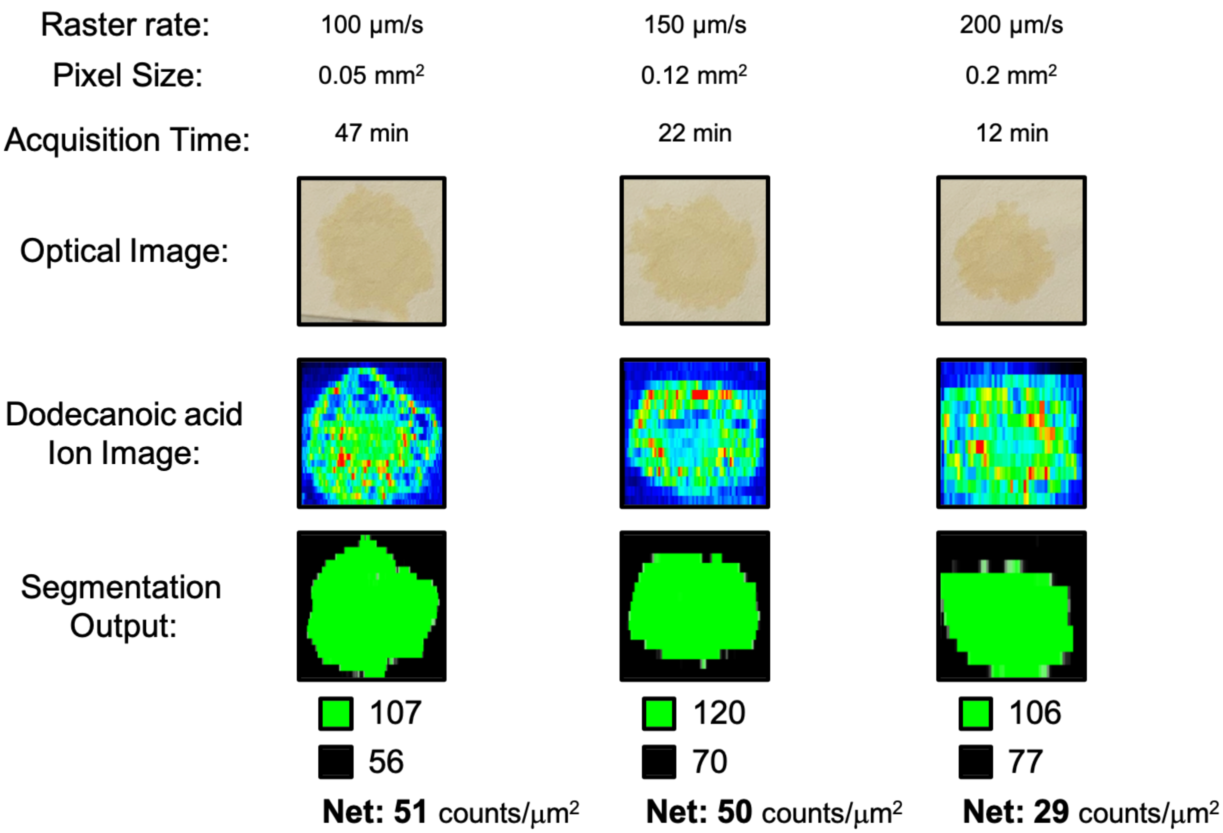


**Supplemental Figure 1:** Evaluation of raster rate on sampling time and dodecanoic acid measurement. Lower raster rates and smaller pixels equate to more resolved images, but longer acquisitions. Increasing raster rate from 100 µm/s to 150 µm/s had a negligible effect on the amount of dodecanoic acid measured from the colony when compared to the background (as defined by unsupervised segmentation). Increasing raster rate to 200 µm/s had an effect on measured amounts of dodecanoic acid.

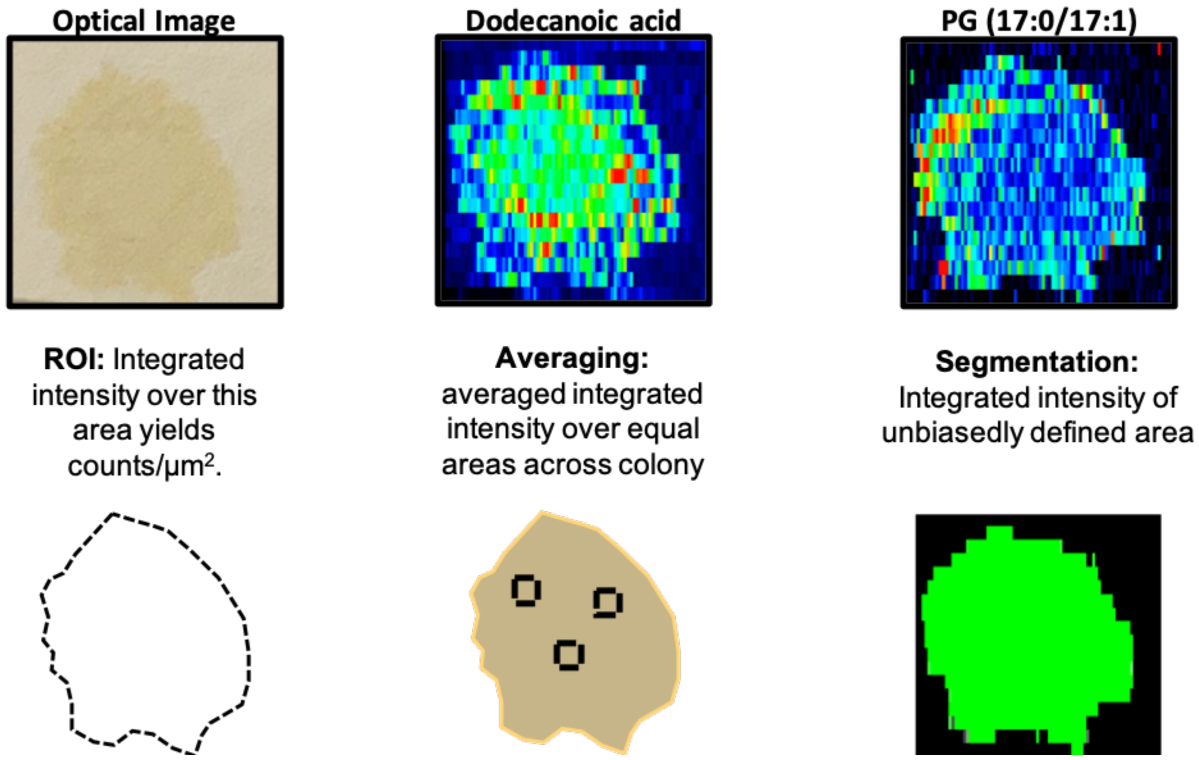


| Data Analysis Method | Average Counts/μm$^2$ (n=3) | Signal-to-noise ratio | Net Counts/μm$^2$ |
|---|---|---|---|
| ROI | 322.3 $\pm$ 11 | 29.3 | 216.7 |
| Averaging | 332.7 $\pm$ 27.3 | 12.2 | 243.8 |
| Segmentation | 333 $\pm$ 15 | 21.8 | 219.3 |

**Supplemental Figure 2:** We compared average counts/μm$^2$, signal-to-noise ratio, and net counts/μm$^2$ of dodecanoic acid from TY05 colonies using the three IMS data analytics methods represented above. Notably, we observe comparable results between segmentation and ROI averaging. While ROI has better performance metrics, it requires user input to define colony areas. Unsupervised segmentation achieves similar output while defining colony areas unbiasedly.

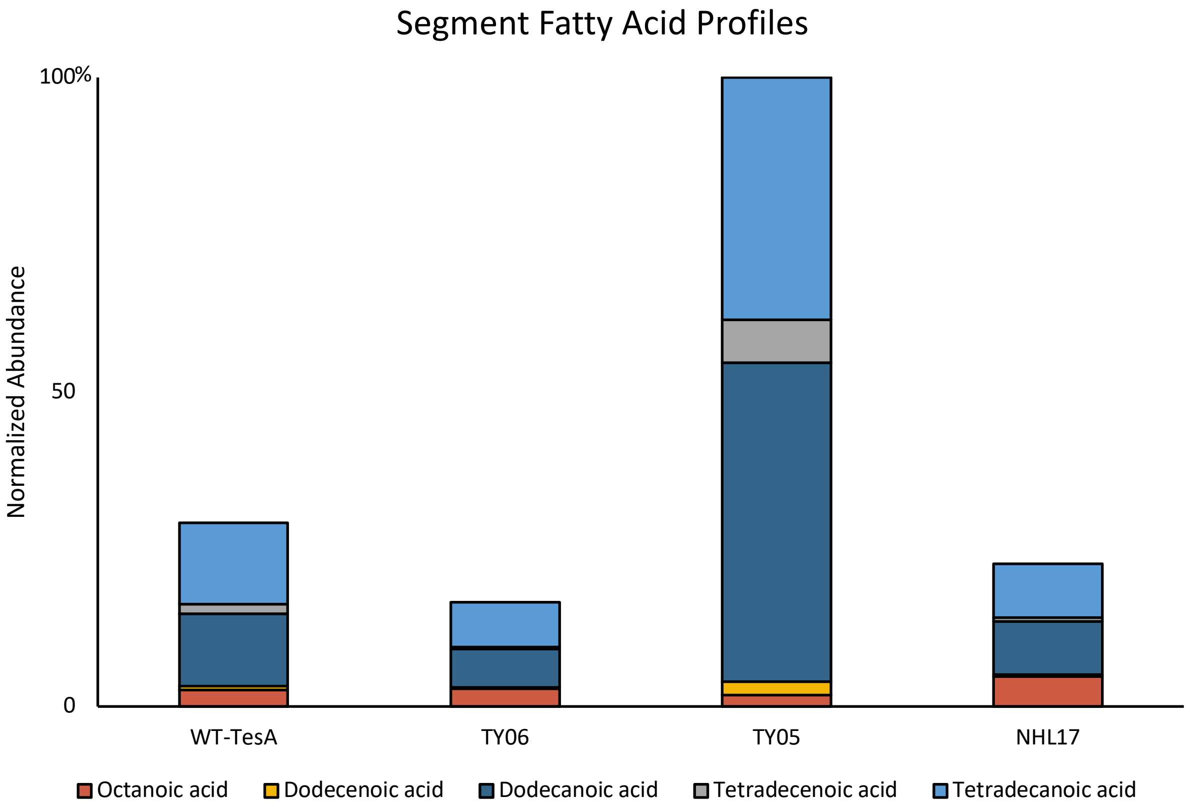


**Supplemental Figure 3:** Segment fatty acid profiles. The IMS areas associated with segments validated via PCR genotyping were integrated to yield comprehensive fatty acid profiles.

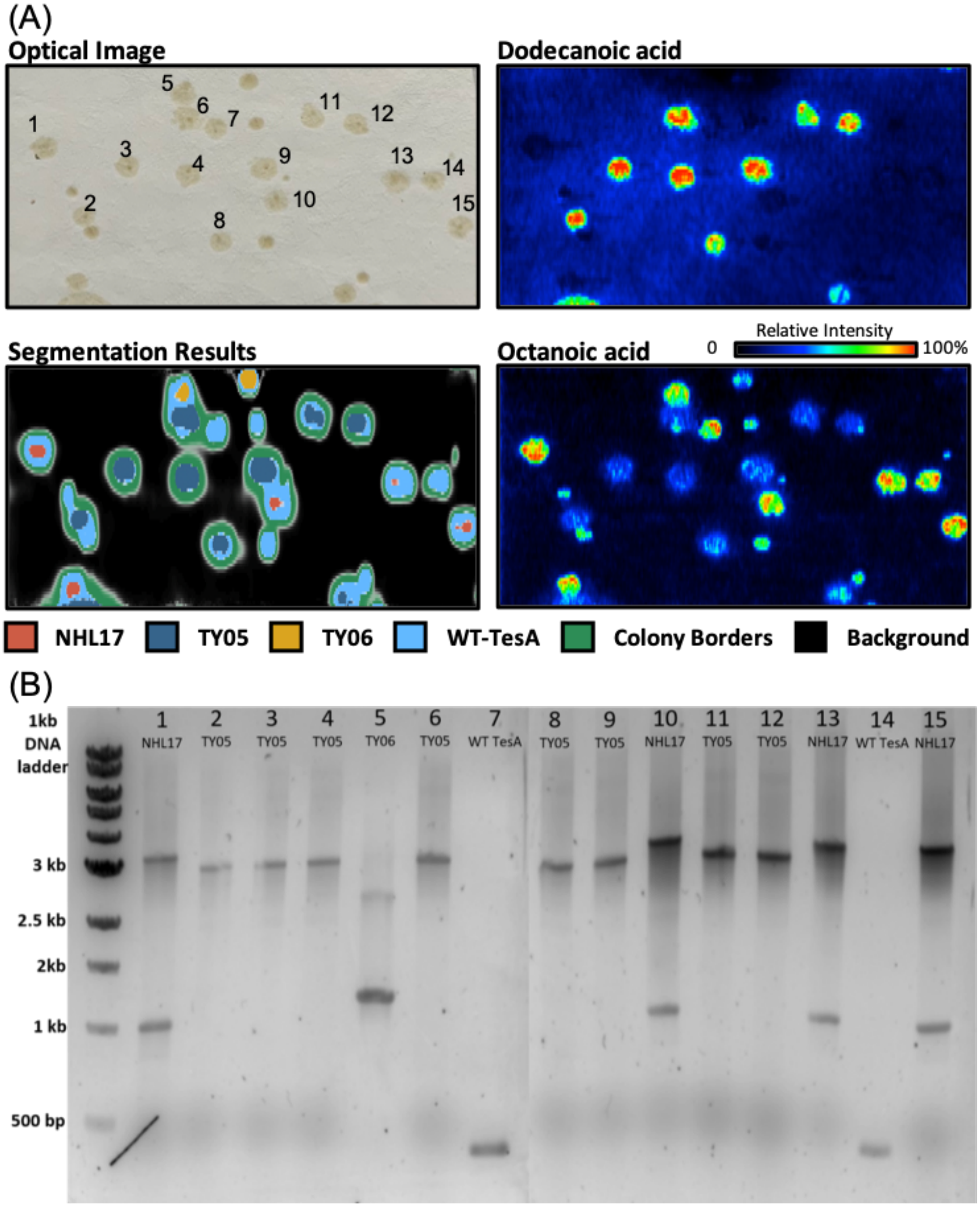


**Supplemental Figure 4:** Confirming unsupervised segmentation results using PCR genotyping. (A) Optical image highlighting colony numbers. Segmentation results highlight the colony areas and identities. Dodecanoic acid and Octanoic acid highlight the primary biosynthetic products contributing to phenotypes. (B) Numbered colonies were streaked on a fresh LB media plate. These replicate colonies were subjected to PCR genotyping instead of original colonies as DESI-IMS may perturb the sample integrity. Colonies that are not numbered formed after this picking process and thus were not analyzed via PCR analysis. PCR primers for each strain are found in supplemental table 1.

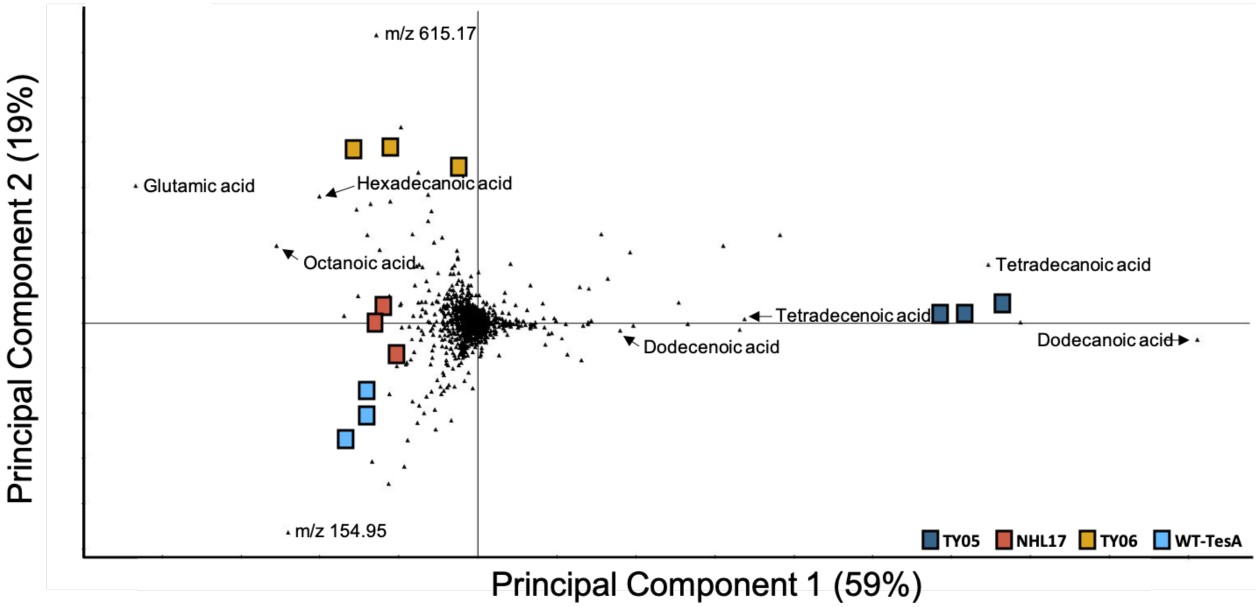


**Supplemental Figure 5:** PCA Loadings plot highlighting features contributing to strain metabolic phenotypes.

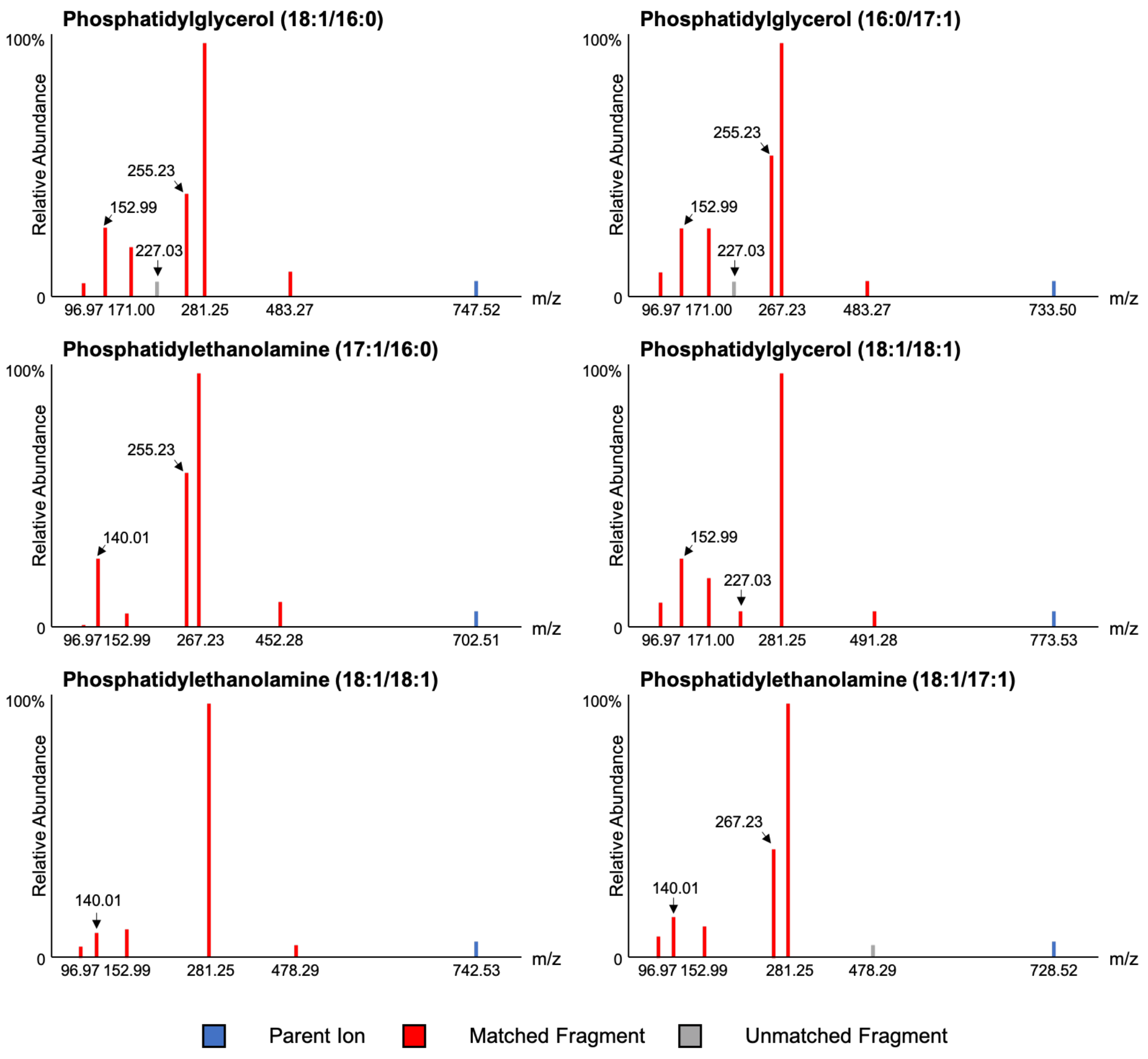


Supplemental figure 6: MS/MS Spectra of Identified Lipids. DESI-MS/MS was performed by directly sampling colonies using the microporous membrane scaffold method. Only features that localized to colony areas were considered as fragment ions. All MS/MS experiments were performed at collision energies of 30 V.

**Table 1.** List of *E.coli* strains, genotype and primer used for their validation.

| Strains | Genotype/construct | Primers | Reference |
|---|---|---|---|
| pBAD18-tesA WT | pBAD18 carrying WT tesA under $P_{BAD}$ control, $Amp^R$ | Forward (TesA-seq-FW) GCCTGCCTTGTTGAATGATAA Reverse(TesA-seq-RV) AATACCGTCATCCTGCATCC | **18:** Grisewood *et al.* 2017 |
| NHL17 | K12 MG1655 ΔaraBAD ΔfadD::trc-CpFatB1.2-M4-287 (C8-specific *Cuphea palustris* FatB1 thioesterase) | Forward (rNHL115) GCATCGTCCGTGGTAATCATTTG Reverse(rNHL93) GCATTTATGCCGATGTGAACGG | **16:** Hernández Lozada *et al.* 2018 |
| TY05 | K-12 MG1655 ΔfadDEAB::trcBTE (acyl-ACP thioesterase from *Umbellularia californica*) | Forward (fadABKO_colPCR_fwd) GGAGTGAATAAGTAACGCATCC Reverse (fadABKO_colPCR_rv) GCTGTCGCGTCTTATCGTGC | **19:** Youngquist *et al.* 2012 |
| TY06 | K-12 MG1655 DfadDEAB::trcBTEH204A | Forward (rNHL229) AGGCAAATTCTGTTTTATCAGACC Reverse (gCRM323) GCACTCCCGTTCTGGATAATG | **19:** Youngquist *et al.* 2012 |
